# Evidence for subsoil specialization in arbuscular mycorrhizal fungi

**DOI:** 10.1101/252544

**Authors:** Moisés A. Sosa-Hernández, Julien Roy, Stefan Hempel, Matthias C. Rillig

## Abstract

Arbuscular mycorrhizal (AM) fungal communities are now known to vary with depth in arable land. Here we use two previously published high-throughput Illumina sequencing data sets, and compare a 52 year long chronosequence of recultivated agriculture fields after a topsoil and subsoil mixing event, with a set of undisturbed topsoil and subsoil samples from a similar field. We show that AM taxa identified as subsoil indicators are exclusively present in early stages of the chronosequence, whereas topsoil indicator taxa can be found across the chronosequence, and that similarities from the chronosequence fields to the subsoil communities decrease with time. Our results provide evidence on the ecological specialization of certain AM fungal taxa to deep soil layers.

## 1. Introduction

Arbuscular mycorrhizal (AM) fungi belong to the monophyletic subphylum Glomeromycotina (Spatafora et al., 2016) and form a symbiotic relationship with most land plants (Brundrett and Tedersoo, 2018). These fungi can increase plant productivity (Lekberg and Koide, 2005), enhance nutrient uptake (Smith and Smith, 2011), promote soil aggregation (Leifheit et al., 2014), boost pathogen protection (Veresoglou and Rillig, 2012), and are therefore considered important factors in agriculture. AM fungal communities in arable land have been characterized both with spore identification techniques (Antunes et al., 2012; Köhl et al., 2014) and molecular methods (e.g. Alguacil et al., 2008; Van Geel et al., 2017) but with few exceptions, existing information is limited to the first 30 cm of the soil profile. Subsoil (i.e. beneath the plough layer) AM fungal communities, however, differ from those in topsoil in diversity, species composition and community structure (Muleta et al., 2008; Oehl et al., 2005; Yang et al., 2010) and even exhibit contrasting patterns of distribution at higher taxonomic levels (Sosa-Hernández et al., 2018). We hypothesize that these differences are caused by Grinellian ecological specialization (Devictor et al., 2010), i. e. top‐ and subsoil represent two different environments to which particular AM taxa have adapted.

A recent study by Roy et al. (2017) used high-throughput Illumina sequencing to analyze AM fungal communities in a series of agricultural fields in western Germany forming a recultivation chronosequence (hereafter referred to as “chronosequence fields”). In short, following mining operations, pits were closed and restored with local soil and after a 3-year period of alfalfa (*Medicago sativa*) cultivation reconverted to conventional agriculture. The restoration was carried out with a mixture of former agricultural soil and loess parent material from various depths. Therefore, we assume that directly after conversion AM fungal communities from different depths experience a community coalescence event (Rillig et al., 2015), i.e. taxa from different depths are mixed in the newly deposited top layers. This event provides excellent opportunity to trace the fate of subsoil-specific AM fungal taxa along the re-cultivation chronosequence, which allows testing our hypothesis of ecological specialization of certain AM fungal taxa to deep soil layers. In a recent study, we characterized AM fungal communities in an agricultural field both in top (10-30cm deep) and subsoil (60-75 cm deep) (Sosa-Hernández et al., 2018), hereafter referred to as “unmixed field”. We identified subsoil and topsoil indicator AM fungal taxa. Here, we traced those taxa along the chronosequence fields. According to our hypothesis, AM fungal taxa identified as subsoil indicators would decrease in abundance in the topsoil along the chronosequence as a function of time since the mixing event occurred, while taxa identified as topsoil indicators will maintain their abundance, and ii) early mixed community would resemble subsoil communities and this similarity would decrease through time.

## 2. Material and methods

### 2.1 Study sites and sequencing

Both study sites are located in the southwest of the state of North Rhine-Westphalia, Germany, and in both soil has been characterized as Haplic Luvisol (FAO, 1998). The chronosequence fields (Roy et al., 2017) consist of a re-cultivation chronosequence after open mining, comprising 10 fields. The newly deposited soil profile is about 2 m deep and consists of a mixture of the previous soil (1 m deep) and loess substrate in a 1:5 ratio. For the first three years after the mixing event fields are covered permanently with alfalfa (hereafter referred to as phase 1), for the two following years barley (*Hordeum vulgare*) was cultivated (hereafter referred to as phase 2) and after the fifth year conventional agriculture was resumed, with a sugar beet (*Beta vulgaris vulgaris* var. *altissima*) - winter wheat (Triticum aestivum) crop rotation (hereafter referred to as phase 3). From these sites five samples per field were taken at a 0-10cm depth, adding up to a total of 50 samples. In the unmixed field nine samples each were taken at depths from 10-30cm and 60-75cm as described in Uksa et al. (2014), adding up to a total of 18 samples. Chicory (*Cichorium intybus*) was grown on this field for the third year.

In both studies AM fungal communities were characterized with primers targeting the large ribosomal subunit LSU including the variable D1-D2 region, using similar protocols (see Roy et al. (2017) and Sosa-Hernández et al. (2018) for details). In short, after DNA extraction PCR was carried out using AM fungal specific primer sets described in Krüger et al. (2009). The product of this amplification was used as a template in a follow up PCR using the general fungal primers LR3 and LR2rev (Hofstetter et al., 2002). Amplicons from the two different studies were sequenced independently but with identical protocols on an Illumina MiSeq platform at the Berlin Center for Genomics in Biodiversity Research (BeGenDiv, Berlin, Germany).

### 2.2 Bioinformatics processing of amplicons sequences

A total of 2,377,171 raw sequences from the chronosequence experiment and 1,876,440 raw sequences from the unmixed experiment were processed separately as follows: Paired-end sequences were merged and quality filtered (maximum error rate of 1) using USEARCH v8.1.1861 (Edgar, 2010). Sequences were dereplicated and singletons were removed. Further quality filtering was performed by aligning those sequences to an AM fungal ribosomal DNA reference database (Krüger et al., 2012) using mothur v.1.38.1 (Schloss et al., 2009), this process also eliminated the primer sequence. Sequences not overlapping the region were discarded.

Quality filtered and dereplicated sequences from the chronosequence experiment (58,686 sequences) and from the unmixed experiment (53,595 sequences) were pooled together and clustered into operational taxonomic units (OTUs) at a 97% similarity level using UPARSE (Edgar, 2013), which includes internal chimera removal. OTU centroids were identified and non-dereplicated filtered sequences from both experiments including previously discarded singletons, were mapped to those OTUs centroids at a 97% similarity level. Various format editing steps such as sequence counting were performed with OBITools 1.2.9 (Boyer et al., 2016).

Taxonomic assignment of the OTUs was carried out using BLAST+ (Camacho et al., 2009) against Glomeromycotina reference sequences published in Krüger et al. (2012) and against the EMBL nucleotide database (Kanz et al., 2005). Alignments below 70% similarity and/or shorter than 300 bp were discarded. Results from both databases were checked for consistency and matches contained in Krüger et al. (2012) were used to assign the OTUs. We decided to favor matches in Krüger et al. (2012) over EMBL, due to the often imprecise description of the match in the latter (e.g. ”soil fungus”, “uncultured Glomeromycota”). When the taxonomic resolution of the match was sufficient, we followed a similar approach to that used in Martínez-García et al. (2015) for SSU sequences, and assigned OTUs with ≥97% similarity match to a species, ≥90% to a genus, ≥80% to a family and ≥70% to the subphylum. In cases with insufficient resolution in the match description, the OTUs were assigned to the closest available taxonomic level. A species level match refers to how confidently we assign a name to our OTU based on known sequences, and does not imply that these OTUs are to be considered equivalent to those species.

### 2.3 Statistical analysis

All subsequent analyses were performed with R version 3.3.1 (R Core Team, 2016). Community analyses were performed with the package “vegan” (Oksanen et al., 2016). Before conduction the statistics, five samples belonging to the 45-year old samples in the chronosequence experiment, were excluded from any subsequent analysis due to very low numbers of AM fungal sequences reads. After this removal, the lowest amount of reads in a sample was determined as 559 and all samples were normalized to this number by random subsampling without replacement with the function “rrarefy”.

Using the sequences retrieved from the unmixed field samples we identified sub and topsoil indicator OTUs using the function multipatt() in the package “indicspecies” (Cáceres and Legendre, 2009) and traced their fate in the the chronosequence since the coalescent event.

Compositional changes between samples were measured with Bray-Curtis (Bray and Curtis, 1957) and Jaccard (Jaccard, 1912) dissimilarities with the function “vegdist” and visualized with a non-metric multidimensional scaling (NMDS) using the function “metaMDS”. Additionally, we compared these Bray-Curtis and Jaccard distances between unmixed topsoil or unmixed subsoil samples to the samples from the chronosequence to test for changes in multivariate distances over time. Comparisons between dissimilarities in different phases were performed with pairwise Mann–Whitney tests with correction for multiple testing, as implemented with the function “pairwise.wilcox.test”.

## 3. Results

After taxonomic assignment and normalization, we identified a total of 136 AM fungal OTUs. Details on the taxonomic assignation of each OTU can be found in **Table S1**. The chronosequence fields yielded a diversity of 123 OTUs and the unmixed fields a diversity of 73 OTUs. Between the two experiments 60 OTUs were shared, representing 44.12% of the total diversity but 93.49% of the reads in “unmixed” fields and 76.53% of the reads in “chronosequence” fields.

We identified three subsoil indicator OTUs (**Table 1**), and we detected two of these subsoil indicator OTUs in topsoil from chronosequence fields with time since the mixing event up to five years (**Fig. 1a**). However, we did not detect these OTUs in chronosequence fields older than five years, neither in the rarefied nor in the non-normalized raw OTU tables. Similarly we identified nine topsoil indicator OTUs (**Table 1**). Those topsoil indicators could be detected in all chronosequence fields and they showed a tendency to increase in relative abundance after the first two years since the mixing event (**Fig. 1b**).

**Fig. 1.**
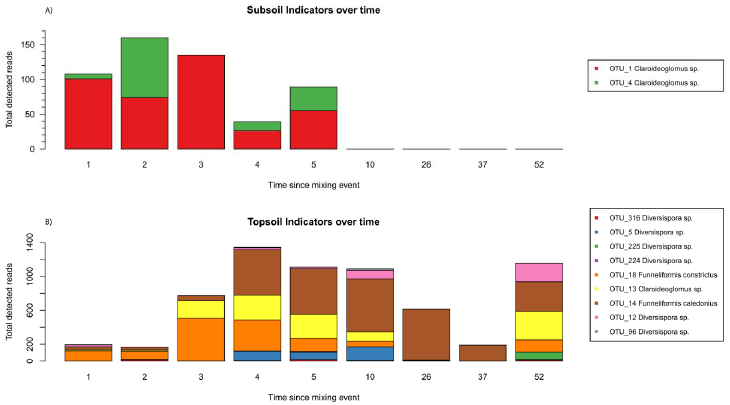
Sub‐ and Topsoil indicators over time. Number of reads detected in the chronosequence fields, for each of the subsoil A) and topsoil B) indicators identified in the unmixed field. Horizontal axis represents the time since the recultivation started, in years. Different indicator OTUs are coded by color.

AM fungal communities in recently restored chronosequence fields (i.e. shortly after the mixing event) are more similar to unmixed subsoil communities, and with increasing time since mixing, chronosequence communities show increasing dissimilarity to unmixed subsoil communities (**Fig. 2**).

**Fig. 2.**
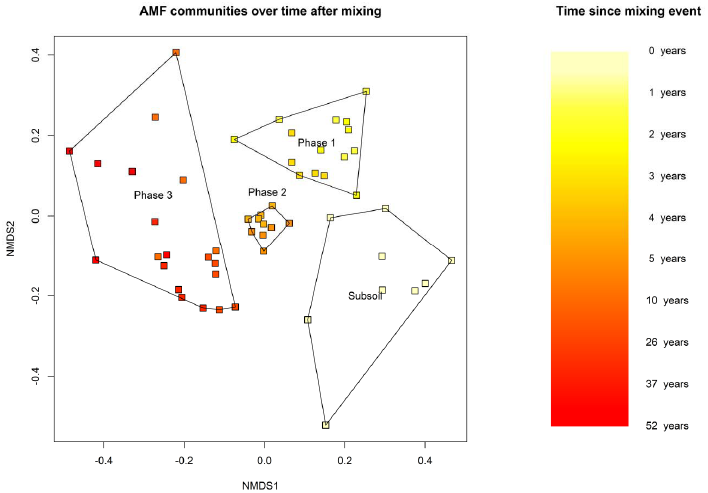
Community ordination of AMF over time. Non-metric multidimensional scaling (NMDS) of a Bray-Curtis pairwise dissimilarity of the AMF communities. The OTU table was rarefied to 559 reads, the minimum amount of reads per sample and includes all chronosequence samples and subsoil samples from the unmixed field. Time since start of the recultivation is coded by color. The polygons encompass all samples from that group. Subsoil = 60–75 cm, n=9. Phase 1: 1-3 years, n=15. Phase 2: 4-5 years, n=10. Phase 3: 10-52 years, n=20.

Bray-Curtis distances from chronosequence fields to the unmixed subsoil samples increase with time, forming two significantly different groups (phase 1 + phase 2, and phase 3; **Fig. 3A**, for statistics see **Table S2**). Analogous results are obtained when considering Jaccard distances (**Fig. S1A,** for statistics see **Table S2**). Bray-Curtis distances to the unmixed topsoil communities follow a unimodal trend with intermediate values in phase1, minimum values in phase 2, and maximum dissimilarity values in phase 3 (**Fig. 3B**, for statistics see **Table S2**). Similarly, Jaccard distances to unmixed topsoil follow a unimodal trend with minimum values in phase 2, but phases 1 and 3 are not significantly different (**Fig. S1B,** for statistics see **Table S2**).

**Fig. 3.**
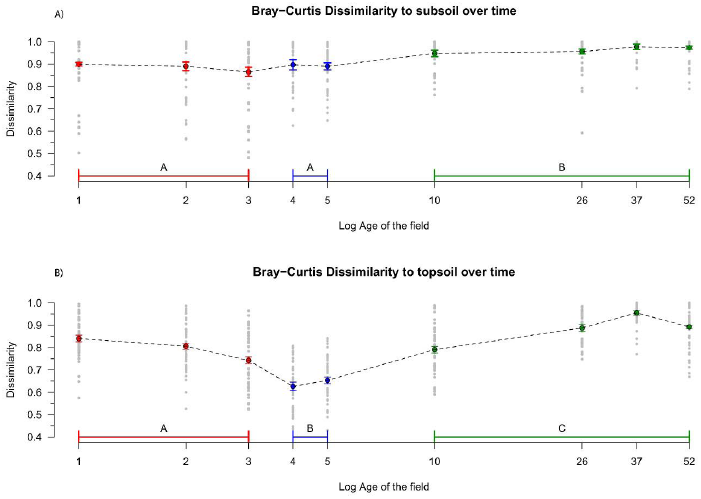
Dissimilarities to Sub‐ and Topsoil over time. Bray-Curtis distances (i.e. dissimilarities in both community composition and relative abundances) between chronosequence fields and A) subsoil communities and B) topsoil communities. Dotted lines link the means, bars represent the standard error. Different phases are coded by color, significant differences between phases are represented by different letters. Details on the statistics are presented in **Table S2**. Phase 1: 1-3 years, n=135. Phase 2: 4-5 year, n=90. Phase 3: 10-52 years, n=180.

## 4. Discussion

We show that i) AM fungal taxa identified as subsoil indicators are present only in young fields (1-3 year since the mixing event), while taxa identified as topsoil indicators are present across the entire chronosequence and ii) early mixed communities from the chronosequence resembled to some extent unmixed subsoil communities and this similarity decreased through time after the mixing event. These results strongly suggest the inability of subsoil-specific AM fungal OTUs to persist in topsoil after a subsoil-topsoil mixing event. Additionally, the detection of topsoil indicators through the entire chronosequence suggests that the observed loss was specific to subsoil phylotypes rather than a generalized diversity loss due to soil treatment during initial deposition or subsequent management.

There are essentially two, not mutually exclusive hypotheses to explain this inability to coexist in the topsoil: abiotic filtering and biotic interactions (Vályi et al., 2016). Possible abiotic filters to subsoil AM fungal taxa in topsoil layers include disturbance in the form of tillage (Kabir, 2005) and greater diurnal and seasonal variations in temperature and moisture (Fierer et al., 2003). Alternatively, possible biotic filters are competitive exclusion by topsoil AM fungal taxa, increased grazing pressure or differential partner selection by the plant due to different nutrient availability. Particularly interesting is the notion that plants might demand different services from AM fungal communities at different depths. By allocating carbon selectively to the desired phylotypes (Werner and Kiers, 2015) plants may shape the observed vertical distribution in AM fungal taxa. It is not clear what the relative importance of abiotic filtering and biotic interactions in driving this species loss is, that is to say, whether subsoil is for this AM taxa a fundamental or a realized niche (Devictor et al., 2010). Equally unknown is whether the subsoil phylotypes established in topsoil of the chronosequence fields and disappeared after a period of time or if they never did establish and the sequences we detect represent dormant inoculum or relic DNA (Carini et al., 2016). We believe that patterns in dissimilarity from “chronosequence” fields to the unmixed topsoil and subsoil communities with time can be interpreted as indirect evidence of the fate of these respective communities across the chronosequence. The slow increase in dissimilarities to unmixed subsoil with time may point at an inactivity and/or slow decline of these OTUs in topsoil, regardless of the host plants or the management. Nonetheless, the observed pattern could as well be explained by the presence and slow decay of relic DNA, as mentioned above. In contrast, the dissimilarities to unmixed topsoil are more responsive to the changes in management in the different phases, suggesting that the members of these communities were active and their populations were part of dynamic turnovers.

Overall, our results support our hypothesis of an ecological specialization of certain AM fungal taxa to deep soil layers. Identifying the specific mechanisms driving the observed patterns will require experimental approaches such as greenhouse reverse transplant experiments or in-vitro competition trials. Nonetheless, our results provide a first snapshot of the outcome of top‐ and subsoil community coalescence events. They show that AM fungal taxa found in subsoils are not able to persist in topsoil layers for longer periods of time. Some deep tillage practices, including deep ploughing or deep mixing, can have positive effects on yield under particular scenarios (Schneider et al., 2017); however, our results suggest that any practice inverting the soil profile has the potential for deleterious effects on AM fungal diversity. Therefore, we suggest that such practices should only be considered as extraordinary measures in soils with root-restricting layers that meet the criteria for potential benefits of deep tillage (Schneider et al., 2017). Whenever possible, subsoiling (i.e. deep ripping) should be preferred over any practice that inverts or mixes the soil profile. With growing awareness of the potential role of AM fungi in sustainable agriculture (Thirkell et al., 2017) acquiring fine-tuned knowledge about the response of particular AM fungal phylotypes to tillage and soil mixing events is crucial if we are to exploit the potential of mycorrhizal technology (Rillig et al., 2016). Caution is needed while handling subsoil AM fungal communities if we are to not irrevocably alter them even before unearthing their secrets and functional potential.

## 5. Acknowledgements

MR acknowledges funding through the Federal Ministry of Education and Research (BMBF) initiative “BonaRes—Soil as a sustainable resource for the bioeconomy” for the projects Soil3 and INPLAMINT. MS-H thanks Carlos Aguilar-Trigueros for numerous and fruitful discussions on this topic.

## References

Alguacil, M. M., Lumini, E., Roldán, A., Salinas-García, J. R., Bonfante, P., and Bianciotto, V. (2008). The impact of tillage practices on arbuscular mycorrhizal fungal diversity in subtropical crops. Ecol. Appl. 18, 527–536. doi:10.1890/07-0521.1.

Antunes, P. M., Lehmann, A., Hart, M. M., Baumecker, M., and Rillig, M. C. (2012). Long-term effects of soil nutrient deficiency on arbuscular mycorrhizal communities. Funct. Ecol. 26, 532–540. doi:10.1111/j.1365-2435.2011.01953.x.

Boyer, F., Mercier, C., Bonin, A., Le Bras, Y., Taberlet, P., and Coissac, E. (2016). obitools□: a unix-inspired software package for DNA metabarcoding. Mol. Ecol. Resour. 16, 176–182. doi:10.1111/1755-0998.12428.

Bray, J. R., and Curtis, J. T. (1957). An Ordination of the Upland Forest Communities of Southern Wisconsin. Ecol. Monogr. 27, 325–349. doi:10.2307/1942268.

Brundrett, M.C., Tedersoo, L., 2018. Evolutionary history of mycorrhizal symbioses and global host plant diversity. New Phytologist. doi:10.1111/nph.14976

Cáceres, M. De, and Legendre, P. (2009). Associations between species and groups of sites: indices and statistical inference. Ecology 90, 3566–3574. doi:10.1890/08-1823.1.

Camacho, C., Coulouris, G., Avagyan, V., Ma, N., Papadopoulos, J., Bealer, K., et al. (2009). BLAST+: architecture and applications. BMC Bioinformatics 10, 421. doi:10.1186/1471-2105-10-421.

Carini, P., Marsden, P. J., Leff, J. W., Morgan, E. E., Strickland, M. S., and Fierer, N. (2016). Relic DNA is abundant in soil and obscures estimates of soil microbial diversity. Nat. Microbiol. 2, 16242. doi:10.1038/nmicrobiol.2016.242.

Devictor, V., Clavel, J., and Julliard, R. (2010). Defining and measuring ecological specialization. 15–25. doi:10.1111/j.1365-2664.2009.01744.x.

Edgar, R. C. (2010). Search and clustering orders of magnitude faster than BLAST. Bioinformatics 26, 2460–2461. doi:10.1093/bioinformatics/btq461.

Edgar, R. C. (2013). UPARSE: highly accurate OTU sequences from microbial amplicon reads. Nat. Methods 10, 996–998. doi:10.1038/nmeth.2604.

Fierer, N., Schimel, J. P., and Holden, P. A. (2003). Variations in microbial community composition through two soil depth profiles. Soil Biol. Biochem. 35, 167–176. doi:10.1016/S0038-0717(02)00251-1.

Van Geel, M., Verbruggen, E., De Beenhouwer, M., van Rennes, G., Lievens, B., and Honnay, O. (2017). High soil phosphorus levels overrule the potential benefits of organic farming on arbuscular mycorrhizal diversity in northern vineyards. Agric. Ecosyst. Environ. 248, 144–152. doi:10.1016/j.agee.2017.07.017.

Hofstetter, V., Clémençon, H., Vilgalys, R., and Moncalvo, J.-M. (2002). Phylogenetic analyses of the Lyophylleae (Agaricales, Basidiomycota) based on nuclear and mitochondrial rDNA sequences. Mycol. Res. 106, 1043–1059. doi:10.1017/S095375620200641X.

Jaccard, P. (1912). The distribution of the flora in the alpine zone. New Phytol. 11, 37–50. doi:10.1111/j.1469-8137.1912.tb05611.x.

Kabir, Z. (2005). Tillage or no-tillage: Impact on mycorrhizae. Can. J. Plant Sci. 85, 23–29. doi:10.4141/P03-160.

Kanz, C., Aldebert, P., Althorpe, N., Baker, W., Baldwin, A., Bates, K., et al. (2005). The EMBL Nucleotide Sequence Database. Nucleic Acids Res. 33, D29–33. doi:10.1093/nar/gki098.

Köhl, L., Oehl, F., and van der Heijden, M. G. A. (2014). Agricultural practices indirectly influence plant productivity and ecosystem services through effects on soil biota. Ecol. Appl. 24, 1842–1853. doi:10.1890/13-1821.1.

Krüger, M., Krüger, C., Walker, C., Stockinger, H., and Schüßler, A. (2012). Phylogenetic reference data for systematics and phylotaxonomy of arbuscular mycorrhizal fungi from phylum to species level. New Phytol. 193, 970–984. doi:10.1111/j.1469-8137.2011.03962.x.

Krüger, M., Stockinger, H., Krüger, C., and Schüssler, A. (2009). DNA-based species level detection of *Glomeromycota*: one PCR primer set for all arbuscular mycorrhizal fungi. New Phytol. 183, 212–23. doi:10.1111/j.1469-8137.2009.02835.x.

Leifheit, E. F., Veresoglou, S. D., Lehmann, A., Morris, E. K., and Rillig, M. C. (2014). Multiple factors influence the role of arbuscular mycorrhizal fungi in soil aggregation-a meta-analysis. Plant Soil 374, 523–537. doi:10.1007/s11104-013-1899-2.

Lekberg, Y., and Koide, R. T. (2005). Is plant performance limited by abundance of arbuscular mycorrhizal fungi? A meta-analysis of studies published between 1988 and 2003. New Phytol. 168, 189–204. doi:10.1111/j.1469-8137.2005.01490.x.

Martínez-García, L. B., Richardson, S. J., Tylianakis, J. M., Peltzer, D. A., and Dickie, I. A. (2015). Host identity is a dominant driver of mycorrhizal fungal community composition during ecosystem development. New Phytol. 205, 1565–1576. doi:10.1111/nph.13226.

Muleta, D., Assefa, F., Nemomissa, S., and Granhall, U. (2008). Distribution of arbuscular mycorrhizal fungi spores in soils of smallholder agroforestry and monocultural coffee systems in southwestern Ethiopia. Biol. Fertil. Soils 44, 653–659. doi:10.1007/s00374-007-0261-3.

Oehl, F., Sieverding, E., Ineichen, K., Ris, E. A., Boller, T., and Wiemken, A. (2005). Community structure of arbuscular mycorrhizal fungi at different soil depths in extensively and intensively managed agroecosystems. New Phytol. 165, 273–283. doi:10.1111/j.1469-8137.2004.01235.x.

Oksanen, J., Blanchet, F. G., Kindt, R., Legendre, P., Minchin, P. R., O’Hara, R. B., et al. (2016). Vegan: community ecology package. doi:10.4135/9781412971874.n145.

R Core Team (2016). R: A language and environment for statistical computing. R Foundation for Statistical Computing, Vienna, Austria. Available at: https://www.rproject.org.

Rillig, M. C., Antonovics, J., Caruso, T., Lehmann, A., Powell, J. R., Veresoglou, S. D., et al. (2015). Interchange of entire communities: Microbial community coalescence. Trends Ecol. Evol. 30, 470–476. doi:10.1016/j.tree.2015.06.004.

Rillig, M. C., Sosa-Hernández, M. A., Roy, J., Aguilar-Trigueros, C. A., Vályi, K., and Lehmann, A. (2016). Towards an Integrated Mycorrhizal Technology: Harnessing Mycorrhiza for Sustainable Intensification in Agriculture. Front. Plant Sci. 7, 1625. doi:10.3389/fpls.2016.01625.

Roy, J., Reichel, R., Brüggemann, N., Hempel, S., and Rillig, M. C. (2017). Succession of arbuscular mycorrhizal fungi along a 52-years agricultural recultivation chronosequence. FEMS Microbiol. Ecol. doi:10.1093/femsec/fix102.

Schloss, P. D., Westcott, S. L., Ryabin, T., Hall, J. R., Hartmann, M., Hollister, E. B., et al. (2009). Introducing mothur: open-source, platform-independent, community-supported software for describing and comparing microbial communities. Appl. Environ. Microbiol. 75, 7537–7541. doi:10.1128/AEM.01541-09.

Schneider, F., Don, A., Hennings, I., Scmittman, O., and Seidel, S. J. (2017). The effect of deep tillage on crop yields – what do we really know? Agric. Ecosyst. Environ. In review, 193–204. doi:10.1016/j.still.2017.07.005.

Smith, S. E., and Smith, F. A. (2011). Roles of arbuscular mycorrhizas in plant nutrition and growth: new paradigms from cellular to ecosystem scales. Annu. Rev. Plant Biol. 62, 227–250. doi:10.1146/annurev-arplant-042110-103846.

Sosa-Hernández, M. A. M. A., Roy, J., Hempel, S., Kautz, T., Köpke, U., Uksa, M., et al. (2018). Subsoil arbuscular mycorrhizal fungal communities in arable soil differ from those in topsoil. Soil Biol. Biochem. 117, 83–86. doi:10.1016/j.soilbio.2017.11.009.

Spatafora, J. W., Chang, Y., Benny, G. L., Lazarus, K., Smith, M. E., Berbee, M. L., et al. (2016). A phylum-level phylogenetic classification of zygomycete fungi based on genome-scale data. Mycologia 108, 1028–1046. doi:10.3852/16-042.

Thirkell, T. J., Charters, M. D., Elliott, A. J., Sait, S. M., and Field, K. J. (2017). Are mycorrhizal fungi our sustainable saviours? Considerations for achieving food security. J. Ecol. 105, 921–929. doi:10.1111/1365-2745.12788.

Uksa, M., Fischer, D., Welzl, G., Kautz, T., Köpke, U., and Schloter, M. (2014). Community structure of prokaryotes and their functional potential in subsoils is more affected by spatial heterogeneity than by temporal variations. Soil Biol. Biochem. 75, 197–201. doi:10.1016/j.soilbio.2014.04.018.

Vályi, K., Mardhiah, U., Rillig, M. C., and Hempel, S. (2016). Community assembly and coexistence in communities of arbuscular mycorrhizal fungi. ISME J. 10, 2341–2351. doi:10.1038/ismej.2016.46.

Veresoglou, S. D., and Rillig, M. C. (2012). Suppression of fungal and nematode plant pathogens through arbuscular mycorrhizal fungi. Biol. Lett. 8, 214–217. doi:10.1098/rsbl.2011.0874.

Werner, G. D. a, and Kiers, E. T. (2015). Partner selection in the mycorrhizal symbiosis/mutualism. New Phytol. 205, 1437–1442. doi:10.1111/nph.13113.

Yang, F. Y., Li, G. Z., Zhang, D. E., Christie, P., Li, X. L., and Gai, J. P. (2010). Geographical and plant genotype effects on the formation of arbuscular mycorrhiza in Avena sativa and Avena nuda at different soil depths. Biol. Fertil. Soils 46, 435–443. doi:10.1007/s00374-010-0450-3.

